# Population origin, body mass, and viral infections influence drone honey bee (*Apis mellifera*) heat tolerance

**DOI:** 10.1101/2024.05.12.593456

**Authors:** Alison McAfee, Bradley N Metz, Patrick Connor, Keana Du, Christopher W Allen, Luis A Frausto, Mark P Swenson, Kylah S Phillips, Madison Julien, Boris Baer, David R Tarpy, Leonard J Foster

## Abstract

Extreme temperatures associated with climate change are expected to impact the physiology and fertility of a variety of insects, including honey bees. Most previous work has focused on female honey bees, and comparatively little research has investigated how heat exposure affects males (drones). To address this gap, we tested how body mass, viral infections, Africanization, and geographic origin (including stocks from Australia, California, and Ukraine as well as diverse locations within British Columbia, Canada) influenced drone and sperm heat tolerance. We found that individual body size was highly influential, with heavier drones being more likely to survive a heat challenge than smaller drones. Drones originating from feral colonies in Southern California (which are enriched for African genetics) were also more likely to survive a heat challenge than drones originating from commercially-supplied Californian stock. We found no association between drone mass and thermal tolerance of sperm over time in an *in vitro* challenge assay, but experimental viral infection decreased the heat tolerance of sperm. Overall, there is ample variation in sperm heat tolerance, with sperm from some groups displaying remarkable heat resilience and sperm from others being highly sensitive, with additional factors influencing heat tolerance of the drones themselves.

## Introduction

Heat waves are expected to increase in frequency and severity as the climate changes [1], threatening the fertility of insects and other animals [2]. Extreme temperatures within the range of what may occur during summer heat waves (35 – 45 °C) can negatively impact the fertility of flour beetles (*Tribolium castaneum*) [3], fruit flies (*Drosophila sp*.) [4], bumble bees (*Bombus sp.*) [5, 6], fruit moths (*Grapholita molesta*) [7], parasitoid wasps (*Anisopteromalus calandrae* and *Aphidius avenae*) [8, 9], and honey bees (*Apis mellifera*) [10–12], among others [2]. Although many insect species are affected by the heat, the impact on bees is particularly concerning due to our reliance on them for agricultural pollination.

Honey bees are the most widely used pollinating species in agricultural systems. Although *A. mellifera* tolerates a wide range of climates, with native populations in Africa, Asia, and Europe, queens (reproductive females) and drones (reproductive males) appear to still be susceptible to heat-induced loss of fertility [10–13]. Queens can survive heat-stress, but the viability of sperm cells stored within their spermathecae diminishes when queens are exposed to temperatures above 38 °C for an excess of 2 hours (h) [10]. Drones, however, are more susceptible to heat-induced mortality [10, 11], and despite being indispensable for reproduction, little is known about the factors governing how heat impacts their survival and fertility.

Two previous studies show that drones are sensitive to heat, but they do not agree on the magnitude of this sensitivity. McAfee et al. reported that 50% of drones survived a 6 h treatment at 42 °C and 60% relative humidity [10], whereas Stürup et al. reported that just 23% of drones survived a shorter (4 h) treatment the same temperature (humidity not reported) [11]. While there are many potential explanations for discrepancies in results between laboratories, including humidity differences, one possibility ― in part motivating this research ― is that perhaps drones derived from different populations have adapted to different climates, and therefore have different temperature tolerance thresholds.

Previous research suggests that there is a genetic component to honey bee temperature tolerance [14], but existing data are limited to workers (non-reproductive females). Workers of a honey bee subspecies originating from Saudi Arabia (*Apis mellifera jemenitica*) are significantly more tolerant to hot temperatures than honey bees originating from Europe (*Apis mellifera carnica*) [14]. Whether this finding extends to reproductive individuals and different populations has not yet been tested, and expanding our knowledge in this area may help predict the outcomes of different genotypes in a changing climate.

A hotter climate may also lead to surprising interactions with diseases, which could, in turn, affect fertility. There is growing evidence that heat-shock proteins are an important part of the antiviral immune response in honey bees [15, 16] and other insects [17], which suggests that there are interactive effects between temperature tolerance and viral infection. McMenamin *et al*. [15] demonstrated this phenomenon by exposing worker honey bees to heat shortly after infecting them with a virus, which improved the bees’ ability to clear the infection. This phenomenon has not been confirmed in drones, despite drones being disproportionately susceptible to viral infection due to preferential parasitism by the *Varroa destructor* mite (a major vector of honey bee viruses) [18] and implications for fertility have not been explored.

Long-term data show that the number of heat-wave days has increased the most in low-latitude (tropical) regions [19]; however, extreme conditions can still occur in temperate climates. During the heat event in Western Canada and the Pacific Northwest spanning June 25 to July 1, 2021, temperatures were sufficiently extreme for beekeepers in the region to observe negative impacts on their colonies [20]. Therefore, beekeepers in high-latitude regions will likely need to begin employing heat-management strategies that have not been historically necessary. Such strategies may include mechanical approaches (*e.g.* year-round hive insulation, shading, *etc.*), which could support colonies over shorter periods of heat exposure. But since pollination necessarily occurs outside the hive and bees will likely face longer periods of heat exposure in the future, in the long term, strategic stock selection and breeding for climate resilience will be required. Such an endeavor will rely on first identifying a relationship between heat tolerance and population origin among commercially viable stocks.

Southern California has also seen a shift toward more extreme temperatures. The region is characterized by a dry, Mediterranean-like climate, and has historically been classified as semi-arid (Köppen climate classification BSk), but more recently it is more characteristic to an arid climate (Köppen climate classification BWh) with hot, dry summers and mild, relatively wet winters. Concerningly, the frequency of heat-waves have rapidly increased since 1950 [21]. For example, a major heat-wave from September 5-7, 2020, in Riverside County caused daytime highs to reach 47 °C [22], and inland counties such as Riverside and San Bernardino are expected to experience higher temperatures during future heat-waves compared to coastal counties, such as Los Angeles [23]. Interestingly, unmanaged (feral) honey bees in Southern California have been genetically characterized and found to contain approximately 25% African ancestry [24]. Since African honey bees survive in hot climates, and unmanaged honey bees in Southern California have been historically exposed to intense heat, we expect bees from this population to possess adaptations to heat and drought stress.

Because of these collective observations, we sought to investigate factors affecting heat tolerance of honey bee drones, including the influence of population origin, body size, and viral infections. In a common garden experiment, we compared heat tolerance (survival) of drones originating from Ukraine, Australia, and California with variable body sizes. In a second common garden experiment, we conducted a similar analysis of drones from feral (hybridized, with some African ancestry) [24] and commercial (managed, with no African ancestry) lineages in California. Finally, we investigated interactions between body mass, viral infection, and fertility of both naturally infected and experimentally infected drones whose sperm cells were subjected to heat challenges. These experiments collectively help build the framework of variables that influence male honey bee heat tolerance.

## Methods

Experiments in this body of work were conducted at three different locations: the University of British Columbia (UBC), University of California Riverside (UCR), and North Carolina State University (NCSU).

### Queen sources and honey bee colonies at UBC

In March 2022, we determined *Varroa* mite levels of overwintered colonies at UBC using the alcohol wash method. We subsequently treated any colonies yielding > 0.5 mites per 100 bees with Formic Pro, according to the manufacturer’s instructions, to ensure minimal interference of mites with drone brood rearing. In April 2022, we purchased commercially available honey bee queens imported to Canada directly from Australia (New South Wales) and Ukraine (Carpathian region). We introduced five queens from each source into overwintered colonies at UBC which had been de-queened the previous day. After a three-day acclimation period, we released the queens through a candy tube. Five other overwintered colonies were already headed by queens imported from a commercial queen supplier in the USA (California) in 2021, bringing the total colonies initially in the experiment to 15. However, due to instances of queen rejection, poor population build-up, or inadequate drone laying, the final data set is made up of drones derived from six colonies (two per geographic origin).

We fed all colonies 50% sugar syrup and 15% pollen patties (Global) on a weekly schedule to encourage population growth and facilitate drone rearing. In May, we confined established queens on drawn drone comb for three days using a single-frame excluder, which confines the queen but allows workers to pass freely. Any eggs the queen laid were allowed to develop *in situ*. Twenty-six days after the queens were caged on drone comb, we returned to the colonies and marked callow or actively emerging drones with a Posca paint pen. Callow (≤ 1 day old) drones are easily recognizable by their light grey appearance. In cases where two colonies were in the same apiary, the drones were marked with different colors so we could identify drifted drones.

Six days after paint-marking drones, we returned to the colonies and collected marked drones into cages made of inverted, clear plastic cups. The cages had ∼100 air holes (1 mm diameter) melted in the sides with a hot metal comb, and the floor of the cage was made of a wire mesh disc that was hot glued on to the rim. Between 20 and 50 drones were held in each cage (two cages per colony), depending on how many could be recaptured, as well as an equal number of workers. Caged bees were held overnight in a 33 °C incubator with access to 50% sugar syrup fed by a dental wick pressed against the mesh floor as well as a 2 mL microfuge tube packed with fondant in the cage (Ambrosia).

### Survival challenge 1 (Californian, Australian, and Ukrainian drones)

The next day, we moved half the cages (one for each colony) to the hot incubator (42 °C, 60% relative humidity). The other cages remained in the 33 °C control incubator. After four hours, dead (immobile) and live (mobile) drones in all cages were recorded. Dead drones were removed with forceps and weighed. Next, the remaining bees were anaesthetized with carbon dioxide (10 min), workers were removed, and the live drones were weighed. All drones were then frozen in 50 mL tubes for later processing.

### Sperm viability challenge

Ten to twelve drones from each colony were used for the sperm viability challenge. We briefly anaesthetized (1 min) the drones with carbon dioxide, then dissected out their seminal vesicles, which we gently ruptured within 100 µl of Buffer D (17 mM D-glucose, 54 mM KCl, 25 mM NaHCO_3_, 83 mM Na_3_C_6_H_5_O_7_) [25] with a plastic pestle. For some colonies, there are sperm viability data but no drone survival data, which occurred when queens produced sufficient drones for viability testing but not the larger number needed for survival.

We diluted 10 µl of the sperm solution into a further 90 µl Buffer D in two separate tubes (one tube for the heat challenge and one control tube). Once all sperm samples were prepared, half the tubes were moved to the same incubator used for the heat survival challenge (42 °C) and half the tubes were moved to the control incubator (33 °C). After four hours, we assessed sperm viability by dual fluorescent staining using SYBR 14 and propidium iodide (Invitrogen™ LIVE/DEAD™ viability/cytotoxicity kit) for both heat-challenged and control samples. Sperm images were taken immediately using a Cellomics HCS (Thermo) microscope at 20x magnification. Once all images were acquired, a colleague not otherwise involved in the study renamed all files with a random character tag so that the stained sperm cells were counted blind.

### British Columbia drone assessments

In July 2022, we received drones from six queen producers located throughout British Columbia, including from the Okanagan (Source A, D, and F), Sunshine Coast (Source B), Fraser Valley (Source C), and Nechako (Source E) regions (**Table 3**). The donors picked approximately sexually mature drones using the “buzz test” (selecting drones that buzzed strongly when pressed on the thorax against the comb). The drones were packed in queen shipping cages (JZ-BZ) with fondant and one drone per cage with five attendant workers. The cages were then shipped in battery boxes, with loose nurse bees and digital temperature loggers, by express courier (1-2 day shipping). All drones were processed two days after being shipped by the donor, even if they arrived after one day, to keep the caging time constant. We conducted the dissections and sperm viability measurements as described above.

### Queen sources and honey bee colonies at UCR

In April and May 2023, commercial bees originated from packages and nucs purchased from Northern California suppliers. Californian feral drones were sourced from rescued swarms (donated by swarm removal specialists in Riverside, Orange, and San Diego counties) and colonies originally of commercial origin, but which bred with local (southern) genetic stock for at least three generations. All colonies were fed *ad libitum* with pollen and sugar syrup. Oxalic acid treatments for *Varroa* mites were used when colonies reached thresholds above 3% using alcohol washes. Treatments were not applied during the drone rearing period.

By June 2023, all colonies had a single frame replaced with drawn drone comb. Colonies were checked weekly to observe egg laying and drone development. Callow drones were marked with a Posca paint pen during weekly checks and different colors were used to track drones to specific marking dates. Drones older than 14 days were removed and placed into 3D-printed plastic holding cages (10 x 6 x 12 cm) with 30 nurse bees from a non-related colony. Water and honey fed *ad libitum*.

### Survival challenge 2

Between 20-30 drones were placed into each cage (1 cage per colony for control and ∼3 cages per colony for heat-shock, depending on drone populations in the colony). After collection, cages were immediately placed into control and heat-shock incubators for 4 hours at 60% relative humidity. The number of live and dead drones were counted in each cage at the end of the treatment. Sample sizes are described in **Table 4**.

### Drone collection and honey bee colonies at NCSU: Temperature-by-time arrays

Honey bee colonies were maintained at the Dix Honey Bee Research Facility in Raleigh, North Carolina and tested on campus at NC State University. Drones sourced from these colonies were used to conduct two sperm viability temperature-by-time arrays: Array 1 (a temperature-by-time array for drones from 6 different colonies and natural variation in viral infections and body mass) and Array 2 (a temperature-by-time array for experimentally infected drones).

For Array 1, a single round of sampling was performed in June 2021. Drones were collected while returning to the nest entrance. The drones were placed into 15 x 15 cm cages of wood and wire mesh and stored briefly in a separate colony until experimental use. The drones were weighed, manually ejaculated [26], then semen was collected into a glass syringe using a Harbo extraction needle [27]. Semen was placed into a 0.25 mL plastic strip tube containing 0.1 mL buffer D [28] and lightly vortexed.

10 µl was immediately removed and diluted 1:10 into buffer containing dyes from the Invitrogen Live/Dead Sperm viability kit (Thermo-Fisher Scientific) to assess baseline sperm count and viability using a Nexcelom Cellometer (Nexcelom Bioscience, LLC). The remaining sperm sample was then aliquoted into strip tubes, placed into a thermocycler, and sperm samples from eight drones each were held at 5 °C, 15 °C, 30 °C, 45 °C, 52.5 °C, and 60 °C. Sperm was repeatedly sampled at 5, 15, 35, 65, 125, and 245 minutes for sperm count and viability. Drone carcasses were then processed via qPCR for incidental pathogen loads of ABPV, BQCV, CBPV, DWV-A, DWV-B, IAPV, and LSV following methods described by Lee et al [29].

For Array 2, two rounds were performed, the first terminating in June 2023 and the second terminating in July 2023. For each round, drones from a single colony were collected from the nest periphery. Drones were collected according to the “buzz test” and caged as in Array 1. 5 mL tube feeders were stocked with 50% sucrose and tap water. Cages also were supplied with a solid feeder stocked with pollen patty made from ground pollen mixed with 50% sucrose. Cages were additionally supplied with 5 g wet honey cappings placed on the cage bottom, which we found to improve drone survival. Liquid food and water were replaced as needed. Each cage was stocked with 100 newly emerged workers and 30 drones. Cages were kept in a dark incubator set to 30 °C and ∼25% relative humidity (ambient for the area).

Drones were either uninjected (control), injected with 1x PBS (sham), or injected with 500 copies/µl Israeli acute paralysis virus (IAPV) in 1x PBS. Injections were conducted with a microinjector with capillary glass needles (Tritech Research) calibrated to deliver approximately 1 µl per injection as assessed by loading 10 µl of 1x PBS and counting the number of injections until the full volume was delivered. A single calibrated needle was used for the whole study where possible, with sham injections preceding the virus injections in all cases to minimize risk of contamination. Needle calibration was checked after every new treatment and after each new needle installation. Drones were injected through the intersegmental membrane between the 5^th^ and 6^th^ abdominal segments, dorsal to the overlap between the tergite and sternite. Any drone that displayed signs of lethargy, injury, or ejaculation after injection was discarded. Drones were kept for three days after injection at which point the survivors were collected for subsequent analyses.

To assess the sperm viability in Array 2, drones were removed from the field facility and returned to the lab where they were prepared for dissection. Drones were dissected following previously described methods [28]. Briefly, live drones were weighed, then reproductive maturity was determined based on mucus gland and seminal vesicle color [30]. The seminal vesicles were homogenized by light mixing with forceps in 100 µl buffer D [25]. Diluted spermatozoa were mixed by briefly vortexing and 10 µl of the spermatozoa sample was immediately diluted 1:10 into buffer containing dyes from the Invitrogen Live/Dead Sperm viability kit (Thermo-Fisher Scientific) to assess baseline sperm count and viability as for Array 1. The remaining sperm sample was then allocated into strip tubes and placed into a thermocycler held at 52.5 °C and repeatedly sampled at 1, 2, and 4 hours. Drone carcasses were then processed via qPCR to confirm the impact of IAPV as well as for additional background viral targets as for Array 1.

### Proteomics analysis

We analyzed a subset of the surviving drones from Survival Challenge 1 to look for patterns of protein expression in relation to drone body size. At the time of conducting the survival challenge, drones were frozen but not stored with unique identifiers after weighing. Due to condensation forming within the tubes after freezing and thawing, wet body mass measurements were no longer indicative of the drone’s original mass at the time of the heat challenge; therefore, we instead recorded thorax width measurements to use as a proxy for mass, as this metric is expected to be stable.

Crude protein was then extracted from the abdomen of each drone using the same protocol as previously described (6 M guanidinium chloride extraction via bead beating homogenization, followed by overnight precipitation in acetone) [31]. All subsequent steps in preparing the samples for mass spectrometry were conducted exactly as previously described (briefly, using urea digestion buffer, reduction/alkylation with DTT/IAA, digestion with trypsin/Lys-C mix, and dilution with 5 volumes of 50 mM ammonium bicarbonate after 4 hours), except: 1) during the C18 STAGE tip cleanup step [32], samples were desalted by washing with 750 µl (3 x 250 µl) Buffer A (0.5% formic acid, 0.5% acetonitrile), and 2) samples were eluted with 250 µl elution buffer (0.5% formic acid, 40% acetonitrile). After reconstitution in Buffer A, sample concentrations were determined using a NanoDrop (A205 nm) and normalized by diluting all samples to peptide concentrations of 20 ng/µl.

A total of 75 ng of digested peptides were injected onto the liquid chromatography (LC) system. The peptides were separated using NanoElute UHPLC system (Bruker Daltonics) with Aurora Series Gen2 (CSI) analytical column (25cm x 75μm 1.6μm FSC C18, with Gen2 nanoZero and CSI fitting; Ion Opticks, Parkville, Victoria, Australia) heated to 50 °C and coupled to a trapped ion mobility-time of flight mass spectrometer (TimsTOF Pro; Bruker Daltonics, Germany) operated in DIA-PASEF mode. The LC gradient and TimsTOF settings were exactly as previously described [33].

### Proteomics data processing

Peptide sequences were identified and quantified using the library-free diaPASEF search option in DIA-NN (v1.8.1) [34]. Default search options were enabled, except: 1) “unrelated runs” was selected, 2) protein inference was conducted based on protein names within the FASTA database, 3) the neural network classifier was set to “double-pass mode,” and 4) up to two missed cleavages were allowed. We used the complete honey bee proteome downloaded from Uniprot (downloaded Feb 24, 2023), with appended protein sequences of all sequenced honey bee viruses and microsporidia, as well as a universal contaminant protein list obtained from Frankenfield *et al.* [35]. The final database had 25,869 entries and is included as part of the publicly available raw data housed on MassIVE (accession: MSV000092842). All protein quantification data, metadata, and statistical results are also available in **Supplementary Data 2.** Protein and peptide identifications were controlled at 1% FDR.

### Statistical analysis

All statistical analyses were conducted in R [36] using the packages nlme and lme4 [37, 38], except proteomics differential expression analysis, which was conducted using the limma package [39].

To compare drone masses between genetic origins, we first confirmed the data were normal with equal variance using a Shapiro test and Levene test, respectively (p > 0.05 for both). We then used a linear model (lm) to analyze the data, using mass as the dependent variable and geographic origin (Australian, Californian, or Carpathian) as the independent variable, and visually confirmed appropriateness of residual distributions.

To determine if drone survival was linked to body mass and genetic origin, we used a generalized linear mixed model (glmer) with survival as a binomial variable (1 = survived, 0 = died), mass and genetic origin as explanatory variables, and colony as a random intercept variable, with a binomial distribution. Only drones which were both exposed to heat and subsequently weighed were analyzed (N = 257 individuals; Table 1). In a separate analysis, we followed the same approach but included the control drones to evaluate the effect of heat. See **Table 1** for sample size information. Drone mass was not recorded for the survival experiment with feral- and commercial-origin drones, so only heat treatment (heat-shock or control) and genetic origin (feral or commercial) were included (as interacting factors). We used tools within the DHARMa package [40] to confirm appropriateness of model fit.

**Table 1.** Sample sizes for Survival Challenge 1.

| Origin | Colony | Treatment | N - total | N - weighed* |
| --- | --- | --- | --- | --- |
| Australia | F1 | Heat | 50 | 35 |
| Australia | F1 | Control | 50 | 0 |
| Australia | F4 | Heat | 50 | 50 |
| Australia | F4 | Control | 50 | 27 |
| Ukraine | F2 | Heat | 50 | 50 |
| Ukraine | F2 | Control | 50 | 43 |
| Ukraine | R1 | Heat | 40 | 40 |
| Ukraine | R1 | Control | 20 | 20 |
| California | R3 | Heat | 40 | 38 |
| California | R3 | Control | 30 | 30 |
| California | R2 | Heat | 48 | 44 |
| California | R2 | Control | 20 | 0 |

\*Only drones receiving the heat treatment were used to identify relationships between mass and survival.

To assess relationships between sperm viability, heat treatments, and genetic origin in the sperm viability challenge, we used a generalized linear mixed model (glmer). Upon visualizing the data and inspecting residual plots, we noted that variance in sperm viability appeared to depend on heat treatment (variance in viability of heat-treated sperm was larger than control sperm). We also noted that the sperm viability data are non-normal, confirmed by a Shapiro test and density plots.

To account for these two issues, we modeled proportion of dead sperm (rather than percent live sperm) using a gamma distribution. The gamma distribution is weighted heavily to smaller numbers and does not permit zeros (which are present in the live sperm parameter, hence using dead sperm instead), which is suitable for our data in this format. Our final model fit proportion of dead sperm against geographic origin (Australian, Californian, or Carpathian) and treatment (heat-shocked or control) as interacting terms, as well as bee as a random intercept variable to account for repeated measures (each drone’s sperm was split into control and heat treatments, and thus these samples are not independent). We reinspected the residual plots and confirmed appropriateness of fit.

To analyze the domestic sperm viability heat challenge data, we followed the same principles as above. We used a generalized linear mixed model (glmer) to model proportion of dead sperm against donor source and heat treatment as explanatory variables and bee as a random variable. As above, we specified a gamma distribution. We were unable to evaluate an interactive effect due to insufficient power.

The sperm viability data from the temperature-by-exposure time arrays (Array 1 and Array 2) had a different distribution compared to the other sperm viability datasets, and was best modelled by applying an arcsine square root transformation to sperm viability proportions. Data from Array 1 was then modelled using a linear mixed effects model (lmer) with transformed sperm viability as the independent variable, a three-way interaction term of temperature (factor; levels: 30 °C, 45 °C, 52.5 °C), exposure time (numeric), and body mass (numeric). Drone was included as a random intercept variable. Although 60 °C was also tested, we removed this group from the analysis as we were unable to design a well-fitting model when including it; however, we still display these data in the results. Arcsine square root-transformed sperm viability data from Array 2 was analyzed with IAPV counts (ln transformed) and time (levels: 0, 1, 2, and 4 h) as interactive terms and drone as a random effect. Data entry and cleanup was performed using the packages broom [41], readxl [42], and packages from the tidyverse [43]. Linear mixed models were performed using the package lme4 [37] and assessed with packages DHARMa [40], car [44], and emmeans [45].

The proteomics data were analyzed by first filtering to remove contaminant sequences, reverse hits, and proteins identified in fewer than 50% of samples. The data were then log2 transformed and analyzed using limma [39] to identify relationships between protein expression and body size (thorax width), heat treatment, and their interaction. The Benjamini-Hochberg method of controlling false discoveries was used (controlled to 5% FDR). To analyze relationships between heat-shock proteins (HSPs) and viral infection, we selected the four HSPs that varied significantly with heat treatment (A0A7M7FYH1, A0A7M7G737, A0A7M7LI93, and A0A7M7LI93; adjusted p < 0.05) and used a linear model to check for correlations with DWV-B polyprotein (G3F401; the viral protein with the highest coverage in the dataset). Because the heat-shock protein expression followed a bimodal distribution (due to the large-magnitude impact of heat-treatment), expression values in heat-treated and control groups were first mean-centered, then correlated with DWV polyprotein abundance.

### Data availability

All data underlying the drone phenotype, survival, and sperm analyses presented here are available in the file Supplementary Data 1. Raw mass spectrometry data, sample metadata, and the search database are housed on the MassIVE archive (www.massive.ucsd.edu; accession: MSV000092842) as well as Supplementary Data 2. Codes associated with statistical analysis are available in Supplementary File 1.

## Results

To determine if drones from different genetic sources had different heat tolerances, we exposed age-matched drones from Australian, Californian, and Ukrainian origins to a fixed temperature (42 °C, 60% relative humidity) and compared their survival rates. In a second, independent common garden experiment, we tested survival of drones from unmanaged origins in Southern California compared to commercial stock produced in Northern California. In the first experiment, survival rates were overall high in both the heat treatments (92.8%; N = 258 of 278) and control groups (99.1%; N = 218 of 220) (**Figure 1a**). While our final sample sizes (88 – 100 individual drones from two colonies per origin) were not sufficient to draw robust comparisons between these sources, we can at least conclude that potential differences here are not large in magnitude. In the second experiment testing feral and commercial Californian drones, we found a significant interaction between origin and heat treatment (**Figure 1b**; χ^2^ = 12.8, p_(interaction)_ = 0.00035). In the first experiment (in which drone mass was recorded) we observed that the dead drones appeared to be biased towards having a small body size, regardless of origin (**Figure 1c and 1d**). We formally tested for this effect and found that smaller drones were significantly more likely to die during the heat exposure (⍰^2^ = 8.63, p = 0.0033) (**Figure 1e**).

**Figure 1.**
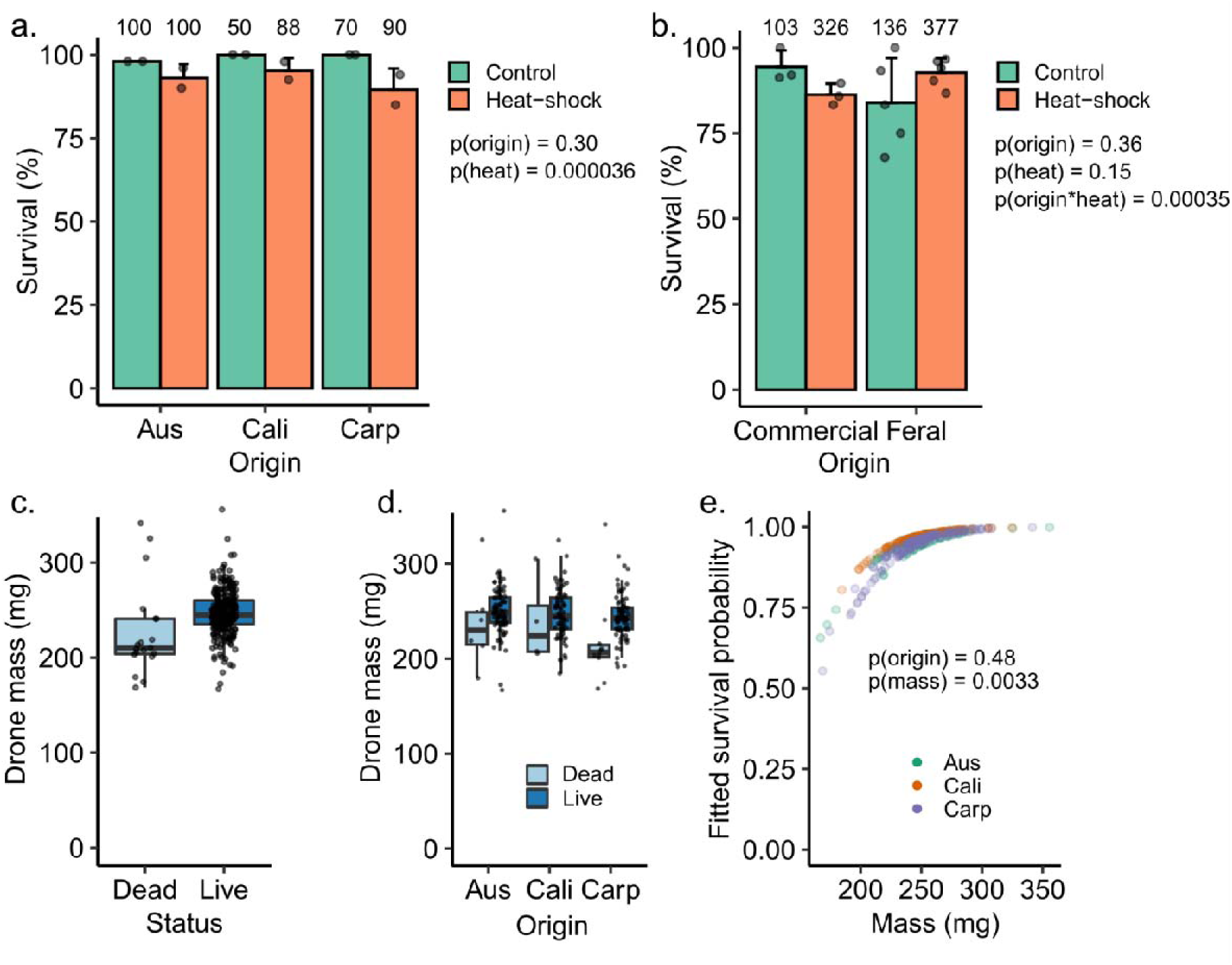
Honey bee drone survival in response to heat. We recorded wet body mass and survival of age-matched drones with different genetic origins (Aus = Australia, Cali = California, Carp = Ukraine) after a 4 h, 42 °C heat challenge. In a separate experiment, we recorded survival of drones from commercial and feral origins in Southern California. A) Drone heat tolerance did not differ among Aus, Cali, and Carp sources. Numbers above bars indicate the total number of drones evaluated in each group. B) Drone heat tolerance interacted significantly with population origin for feral and commercial drones from California. Panel C and D illustrate the data underlying panel E, showing dead and live drone masses together (C) and separated by origin (D). E) Fitted survival probabilities were extracted from the generalized linear model and plotted against wet mass taken immediately after the heat challenge.

To determine if larger drones exhibited a different molecular response to heat than smaller drones, we sampled a subset of heat-treated and control drones from the survival challenge and analyzed proteins extracted from their abdomens by shot-gun proteomics. We tested 45 heat-challenged and 45 control drones, which resulted in 4,414 protein groups identified, 3,887 were quantified (identified in > 50% of samples), and 1,586 were differentially expressed between heat-shocked and control groups at 1% FDR (Benjamini-Hochberg correction), indicating a strong effect of heat (**Figure 2a**).

**Figure 2.**
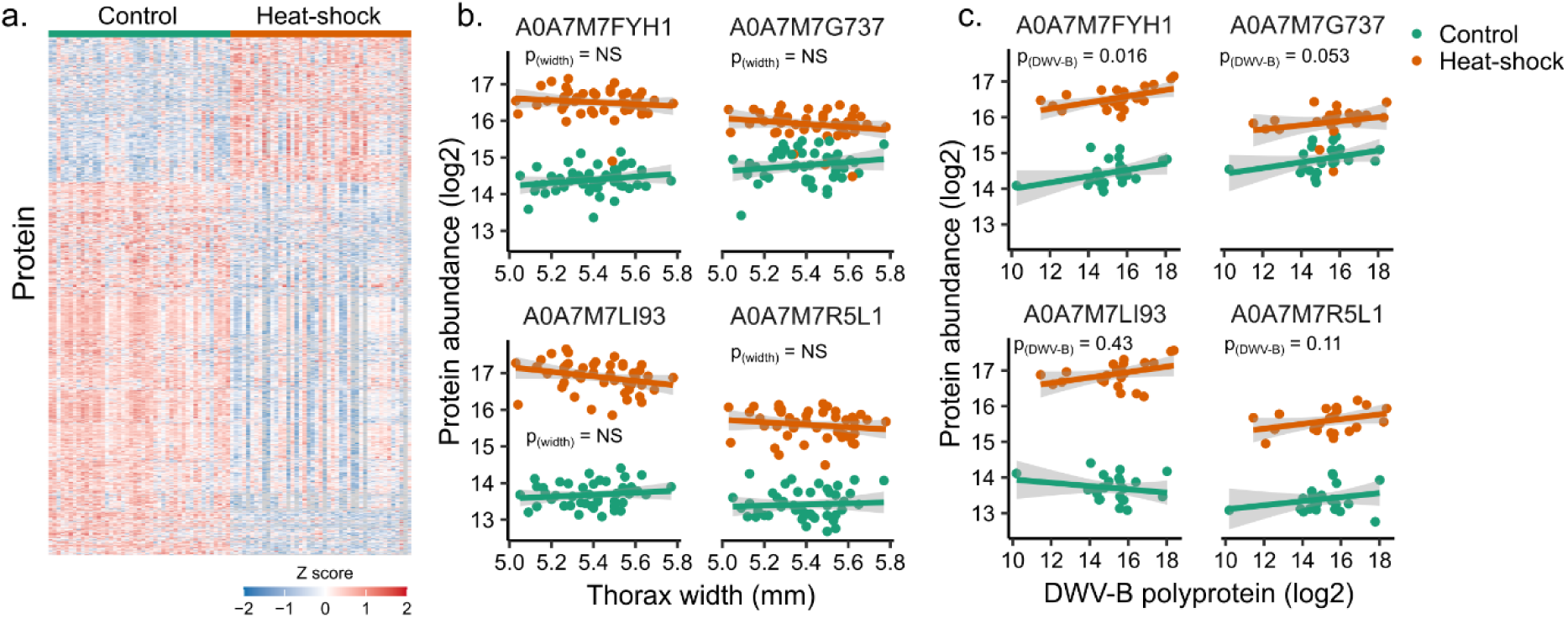
Small heat-shock proteins are induced with heat-shock but do not correlate with drone size. Protein abundances were determined by shot-gun proteomics analysis of 45 heat-shocked and 45 control samples. A) 1,586 proteins (shown) were differentially expressed at 1% FDR (Benjamini-Hochberg) and 2,050 were differentially expressed at 5% FDR. B) No proteins, including these four sHSPs (A0A7M7FYH1, A0A7M7G737, A0A7M7LI93, and A0A7M7R5L1), correlated with drone size, either as a fixed effect or part of an interaction term (limma, adjusted p values all > 0.05). C) The four sHSPs were evaluated individually to test for relationships with DWV-B. Protein abundances were mean-centered ahead of statistical testing. One sHSP (A0A7M7FYH1) significantly correlated with DWV-B polyprotein abundance (F = 6.3, df = 1, p = 0.016).

We were specifically interested in expression patterns of small heat-shock proteins (sHSPs), as these have been previously associated with heat treatments in female bees [10, 15, 46] and their expression patterns may provide insight into the relationship between body mass and survival. We hypothesized that either larger drones have higher baseline levels of sHSPs than smaller drones (priming them to tolerate acute heat stress), or that larger drones may not heat up as fast as smaller drones and thus have a lower magnitude induction of sHSP expression after the heat challenge. We identified four sHSPs in our dataset, and, while all show a strong relationship with heat treatment, no relationships with drone size were identified (**Figure 2b**). The second reason we were interested in sHSPs is because some evidence suggests that some of these proteins are antiviral immune effectors in female honey bees [15, 16], but they have not been investigated in male honey bees. Here, we found that sHSP A0A7M7FYH1 positively correlates with natural DWV-B polyprotein abundance (F = 6.3, df = 1, p = 0.016) and another sHSP, A0A7M7G737, displayed a marginally non-significant relationship (F = 4.0, df = 1, p = 0.053) (**Figure 2c**).

To test if sperm from drones with different origins responded to heat differently, we exposed extracted sperm cells to heat and compared sperm viability between treatments and sources. As predicted, heat significantly reduced sperm viability (**Figure 3a**; ⍰^2^ = 135.2, df = 1, p < 2×10^-16^), but we found no relationship between sperm survival and stock origin (⍰^2^ = 2.52, df = 2, p = 0.28) nor an interaction between treatment and origin (⍰^2^ = 1.55, df = 2, p = 0.46). Drones sourced from domestic colonies (colonies propagated locally at several different queen production operations within British Columbia), however, show a strong effect of local origin on sperm viability (⍰^2^ = 92.7, df = 5, p < 2×10^-16^) and an interactive effect with heat (⍰^2^ = 23.2, df = 5, p = 0.00031) (**Figure 3b**). We were unfortunately not able to evaluate sperm viability of drones sourced from feral and commercial origins in a comparable way, and this remains an important avenue to explore in the future.

**Figure 3.**
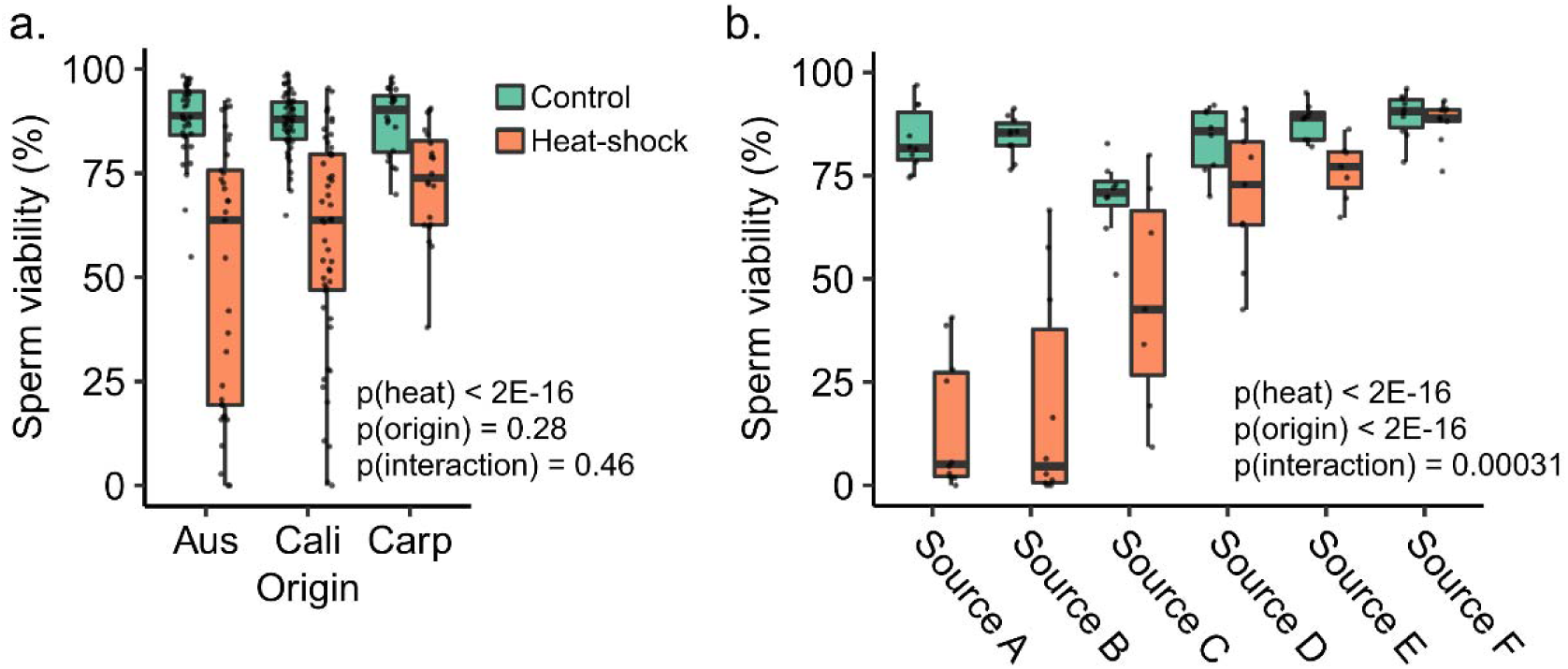
Sperm viability of different genetic stocks and local colonies in response to heat. Sperm cells extracted from seminal vesicles were split into two samples and exposed to heat (42 °C) or control (33 °C) treatments for four hours, followed by viability analysis using dual fluorescent staining. Boxes represent the interquartile range, and whiskers span 1.5 times the interquartile range. The bar represents the median. Bee as a random variable in the generalized linear mixed model. A) Genetic stock comparison. Aus = Australia, Cali = California, Carp = Ukraine. Treatment (heat-shock or control), origin (Aus, Cali, or Carp), and their interaction were included in the linear mixed model, with bee as a random variable (modelling proportion dead sperm, gamma distribution. B) Sperm from drones originating from six different sources throughout British Columbia were tested in the same manner as (A).

Among drones sourced from colonies within British Columbia, there was outstanding variation in sperm temperature tolerance (**Figure 3b**). Such variation could be due to influences of body mass or viral infections that were unaccounted for. To test for relationships between body mass, underlying viral infections, and sperm heat sensitivity, in a separate experiment we recorded wet body mass for N = 48 drones, tested them for a panel of seven honey bee viruses (see Methods), extracted their sperm, and subject them to an *in vitro* array of temperatures and exposure durations.

As expected, we found significant main and interactive effects of exposure time and temperature; however, we found no three-way interaction between time, temperature, and body mass (**Figure 4**). We therefore removed body mass from the interactive term to increase power, but also failed to identify a main effect of body mass on sperm viability (χ^2^_(Mass)_ = 2.37, df = 1, p = 0.12; see **Figure 4** for additional statistics). Natural prevalence of viral infection was not evenly distributed enough among our treatment groups to adequately evaluate interactions between viral infection, temperature, and time, but we did identify a significant effect of the presence of DWV-B in untreated (time = 0) samples, which was associated with higher sperm viability (**Supplemental Figure 1**).

**Figure 4.**
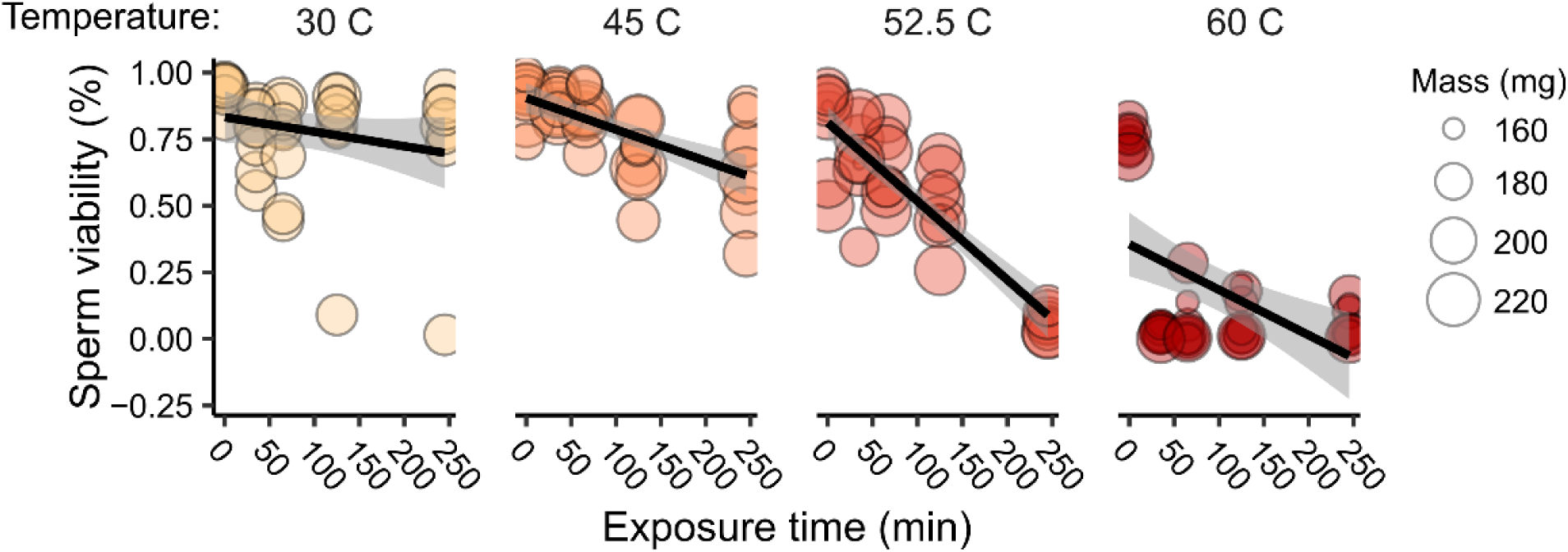
No relationship between body mass and sperm sensitivity to heat. We sampled sperm from 48 drones belonging to six colonies and challenged the sperm to an array of temperatures and exposure durations. We found significant main effects and interactive effects of heat and exposure duration on sperm viability (linear mixed model on arcsine square root-transformed viability proportions; χ^2^_(Temperature)_ = 651, p < 0.0001; χ^2^_(Time)_ = 534, p < 0.0001; χ^2^_(Temperature*Time)_ = 635, p < 0.0001), but no effects of drone mass (χ^2^_(Mass)_ = 2.37, p = 0.12).

These observations, combined with the positive association we observed between sHSP A0A7M7FYH1 and DWV-B earlier (**Figure 2c**), led us to hypothesize that viral infection could paradoxically be associated with higher sperm viability indirectly via antiviral heat-shock proteins [15, 47]. If this is the case, we would expect experimentally infected drones to yield sperm that are also more resilient to heat, as their seminal fluid (and possibly sperm cells) could be primed with higher levels of heat-shock proteins than uninfected drones. To test this, we assessed sperm viability of control, sham-infected, and IAPV-infected drones (n = 9) in response to heat (52.5 °C) over time (0, 1, 2, and 4 h). IAPV was used because there is no evidence for antiviral HSP activity being specific to certain viruses, and the symptomatic nature of IAPV can be an asset for visually determining success of experimental infections.

We confirmed that IAPV copies varied significantly by group (F= 18.85, p<0.0001), with levels in the infected group higher than the sham group by about three orders of magnitude (**Figure 5a**). However, we also observed higher infection levels in the sham group relative to the uninjected drones, which points to some level of viral transfer between groups. IAPV was thus best accounted for in our statistical model for sperm viability as a continuous rather than categorical variable. Applying this approach, we found that IAPV (χ^2^ = 12.35, p=0.0004) and time (χ^2^= 310.49, p<0.0001) significantly and negatively affect sperm viability. These variables also significantly interact (χ^2^= 22.32, p<0.0001) with sperm viability of drones with higher viral loads decreasing faster than control drones (**Figure 5b**).

**Figure 5.**
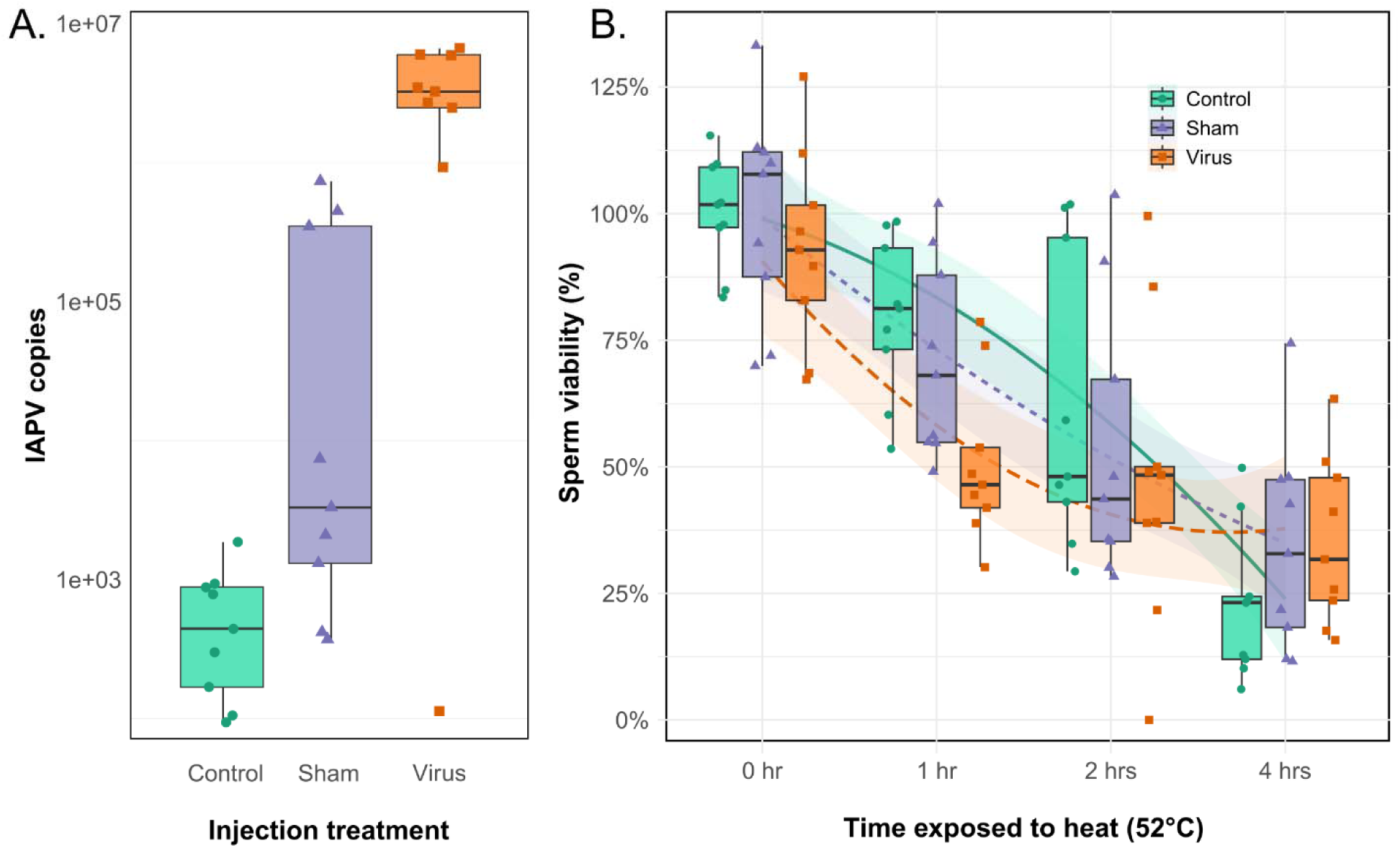
Negative relationship between IAPV infection and sperm viability. We extracted semen from control, sham-injected (saline injection), and IAPV-injected (500 copies of infectious IAPV) drones (n = 9 each) three days after treatment, and exposed sperm to heat for 0, 1, 2, or 4 hours. Boxes represent the interquartile range, and whiskers span 1.5 times the interquartile range. Bars represent the median. A) IAPV copies in the three experimental groups (control, sham, and virus). B) IAPV (χ^2^ = 12.35, p=0.0004), time (χ^2^= 310.49, p<0.0001), and their interaction (χ^2^= 22.32, p<0.0001) all significantly and negatively predict sperm viability.

## Discussion

In an effort to understand what traits contribute to heat resilience of honey bee drones and their mating potential, we aimed to investigate how genetic, physical, and pathogenic factors influence drone heat tolerance. Our main findings are that: 1) heavier drones were more likely to survive an acute heat challenge, regardless of their genetic background (**Figure 1**); 2) drones from feral colonies in Southern California were more likely to survive a heat challenge than their commercial counterparts (**Figure 1**); 3) sperm from heavier drones were not more heat tolerant than sperm from smaller drones (**Figure 3**); 4) sperm from drones with different geographic origins had highly variable levels of heat tolerance (**Figure 4**); and 5) viral infections reduced sperm heat tolerance (**Figure 5**).

The finding that heavier drones are more resilient to heat at first appears to be at odds with the observation that warming temperatures favor arthropods with smaller body sizes [48]. This discrepancy has three potential explanations: 1) since the drones participating in this experiment were from managed stocks, they were under reduced natural selection, and may not show the same heat tolerance trends as wild populations, 2) since we used acute, short-term exposures (which are more biologically relevant in our study system) rather than the longer term, chronic exposures that wild animals also experience, our results reflect a slightly different characteristic, and 3) body mass is influenced by hydration and insulating fat reserves, whereas other size metrics may not be. Without knowing more about the specific selection and management programs used by the commercial suppliers of these drones, the first explanation is difficult to assess. However, we can provide some speculation regarding the second explanation. Larger-bodied individuals both have higher intrinsic heat production and have a lower surface area-to-body mass ratio, which impairs cooling. But in our short-term heat exposures (the longest of which was 4 h), we suspect that larger body masses are favourable because it takes longer for external conditions to raise the core temperature of large drones than small drones. Thus, we propose that the defining factor in the short term is heating time, not cooling capacity. The time scale of exposure that we used is especially relevant for drones, which should only experience extreme conditions for only short spurts during mating flights or periods of maximum sun exposure of the colony (which, unless under exceptionally extreme conditions, is normally well-thermoregulated). This explanation may be acting in combination with the third point, that body mass is influenced by hydration and fat reserves, both of which are expected to reduce the rate of core heating.

We anticipated seeing evidence of the size-dependent heating delay in the proteomics experiment, where we predicted larger bodied drones to exhibit lower HSP expression (indicative of being under reduced stress), but no relationship between HSPs and body size was observed (**Figure 2**). Since this analysis used thorax width rather than body mass as the “size” metric, it is possible that a relationship was obscured because thorax width is an example of a metric that does not take into account variation in hydration and fat reserves. Indeed, this observation lends some credence to the seemingly contradictory positive relationship between heat tolerance and size discussed above. In addition, this analysis may have been more enlightening if a purely internal organ, such as the seminal vesicles, was used instead of whole abdomens, as we would expect these to be more specifically protected from heat. These are both areas which require further experimentation in the future.

We predicted that drones from different geographic or genetic origins may show some indication of differential heat tolerance, either in terms of individual survival or sperm survival. In some cases, this was borne out, and others not. Our first common garden experiment with drones originating from Australia, Northern California, and Ukraine did not yield patterns linked to origin, but these comparisons would benefit from including a larger number of source colonies. Here, our data are limited to drones from two different colonies per source for the survival tests (**Table 1**), and between two and five different colonies for the sperm viability tests (**Table 2**). In addition, an analysis of unmanaged colonies in these regions may be more likely to reveal differences. Perhaps most interestingly, in the second common garden experiment (which did test managed and unmanaged colonies), we found that drones from different origins within California did have different patterns of survival, with unmanaged-origin drones being more likely to survive the heat challenge than commercial-origin drones. This is exciting because the experiment was conducted in Southern California, where feral colonies tend to have a greater proportion of African ancestry than the drones from commercial colonies, which originate from Northern California (outside the Africanized zone) [24]. We speculate that colonies surviving without management in Southern California are under continuous natural selection for heat tolerance. Unfortunately, we do not have corresponding sperm viability for these groups that is comparable to the sperm viability data presented throughout this body of work, but this is an area of investigation that we are actively pursuing.

**Table 2.** Sample sizes for global origin sperm analysis.

| Source | Colony | Group | N |
| --- | --- | --- | --- |
| Australia | F1 | Heat | 11 |
| Australia | F1 | Control | 11 |
| Australia | F4 | Heat | 11 |
| Australia | F4 | Control | 12 |
| Australia | F5 | Heat | 11 |
| Australia | F5 | Control | 12 |
| Ukraine | F2 | Heat | 10 |
| Ukraine | F2 | Control | 10 |
| Ukraine | R1 | Heat | 10 |
| Ukraine | R1 | Control | 10 |
| California | S7 | Heat | 10 |
| California | S7 | Control | 10 |
| California | S4 | Heat | 10 |
| California | S4 | Control | 10 |
| California | S2 | Heat | 10 |
| California | S2 | Control | 10 |
| California | R3 | Heat | 10 |
| California | R3 | Control | 10 |
| California | R2 | Heat | 10 |
| California | R2 | Control | 10 |

**Table 3.** Sample sizes for domestic drone sperm analysis.

| Source | BC region | Group | N |
| --- | --- | --- | --- |
| A | Okanagan | Heat-shocked | 7 |
| A | Okanagan | Control | 9 |
| B | Sunshine Coast | Heat-shocked | 10 |
| B | Sunshine Coast | Control | 10 |
| C | Fraser Valley | Heat-shocked | 10 |
| C | Fraser Valley | Control | 10 |
| D | Okanagan | Heat-shocked | 7 |
| D | Okanagan | Control | 8 |
| E | Nechako | Heat-shocked | 9 |
| E | Nechako | Control | 10 |
| F | Okanagan | Heat-shocked | 9 |
| F | Okanagan | Control | 8 |

**Table 4.** Sample sizes for Survival Challenge 2.

| Origin | Source | Colony | Treatment | N - total |
| --- | --- | --- | --- | --- |
| California - commercial | Package | 402 | Heat | 90 |
| California - commercial | Package | 402 | Control | 30 |
| California - commercial | Package | 408 | Heat | 116 |
| California - commercial | Package | 408 | Control | 23 |
| California - commercial | Nuc | R01 | Heat | 120 |
| California - commercial | Nuc | R01 | Control | 50 |
| California - feral | Swarm | DHS1000 | Heat | 30 |
| California - feral | Swarm | DHS1000 | Control | 28 |
| California - feral | Swarm | R1001 | Heat | 90 |
| California - feral | Swarm | R1001 | Control | 30 |
| California - feral | Swarm | RT1002 | Heat | 25 |
| California - feral | Swarm | RT1002 | Control | 20 |
| California - feral | Multi-generation requeening | 3282 | Heat | 114 |
| California - feral | Multi-generation requeening | 3282 | Control | 28 |
| California - feral | Multi-generation requeening | 4240 | Heat | 118 |
| California - feral | Multi-generation requeening | 4240 | Control | 30 |

We did, however, compare sperm heat tolerance among drones from alternate sources. The sperm viability data from drones throughout diverse regions of British Columbia showed remarkable variation, prompting us to investigate other potential explanatory variables. We were interested in investigating interactive effects of viral infection, in particular, because of three key observations. First, we observed that drones with naturally acquired DWV-B infections had, surprisingly, higher sperm viability than drones without DWV-B infection. Second, our previous research identified a positive relationship between at least one HSP and stored sperm viability [49], and we show a positive relationship here between a different HSP and DWV-B polyprotein abundance. Third, McMenamin et al. have shown that HSPs have antiviral effects [15]; therefore, we hypothesized that a potential mechanism explaining these collective observations could lie in the dual role of HSPs as both antiviral and abiotic stress-mitigating agents.

Although one might expect a viral infection to impair fitness of the individual, we reasoned that perhaps the elevation of HSPs stimulated by the virus could protect sperm from subsequent abiotic stress. However, when we tested this hypothesis by infecting adult drones, then subjecting their sperm to heat stress, we found the opposite — sperm from infected drones were less resilient to heat than sperm from control drones. We have yet to reconcile this observation with the positive correlation between infection and sperm viability we initially found, but it is possible that the disagreement is due to timing of infection (duration of infection before sperm sampling), pathogenicity of the virus (IAPV vs. DWV-B), life stage at which the drone became infected (infections acquired before, during, or after spermatogenesis may produce different outcomes), or a potential consequence of cellular resource limitation in drones that we have yet to deconvolute.

## Conclusion

Here, we report a broad investigation into factors affecting drone heat tolerance, in terms of survival and fertility. Our main findings are that body mass is a significant predictor of drone survival under heat-challenge conditions, that differences in adult survival and sperm heat tolerance exist among drones from different origins, and that adult-acquired viral infection increases sperm sensitivity to heat. This work contributes to building the framework for understanding how drone honey bees and their fertility may be affected under hotter conditions that are already increasing in frequency as the climate changes.

## Acknowledgements

We would like to acknowledge the BC Bee Breeders’ Association, the Long Beach Beekeepers, and the Beekeepers Association of Southern California for their generous assistance and support to UBC and UCR researchers during this project. We would also like to acknowledge the UBC proteomics core facility team – Jason Rogalski, Renata Moravcova, and Jeanne Yuan – for running the mass spectrometry samples, instrument maintenance, and technical expertise. We thank Jennifer Keller for beekeeping support at NCSU, as well as Erin McDermott and Lauren Paturzo for technical training on the pathogen testing.

## Author contributions

AM wrote the first draft of the manuscript, with assistance from BNM and CWA. All authors edited and approved the final version of the manuscript. AM conducted the common garden survival experiment 1, and CWA, LAF, MPS, KSP, and MJ conducted the common garden survival experiment 2. BNM, PC, and KD conducted work related to sperm viability array 1 and array 2. BNM conducted statistical analysis and generated figures for data linked to array 1 and array 2, and AM conducted all other statistical analysis and figure generation. LJF provided funding and infrastructure to support experiments conducted at UBC, BB provided funding and infrastructure to support experiments conducted at UCR, and DRT provided funding and infrastructure to support experiments conducted in NCSU.

## Funding

The mass spectrometry infrastructure (LJF) was supported by the Canada Foundation for Innovation and the BC Knowledge Development Fund. Proteomics operations (LJF) were supported by a Genome Canada/Genome BC project (264PRO). AM’s work was supported in part by a L’Oreal For Women in Science Fellowship, MITACS, and a grant from Project *Apis m.* PC and DU’s work was supported by the BeeMORE REEU program (USDA-NIFA 2021-67037-34626) and BNM was supported by a grant from the US Army Research Office (W911NF1920306) and a grant from the USDA-NIFA (2221-0507). The work conducted in California (CWA, LAF, MPS, KSP, MJ, and BB) was supported by a Multi Campus Research Initiative offered by the University of California Office of the President and OASIS support from the University of California Riverside.

## Notes

### Competing Interest Statement

The authors have declared no competing interest.

