## Supplementary Figure 1 for "Population origin, body mass, and viral infections influence drone honey bee (*Apis mellifera*) heat tolerance"

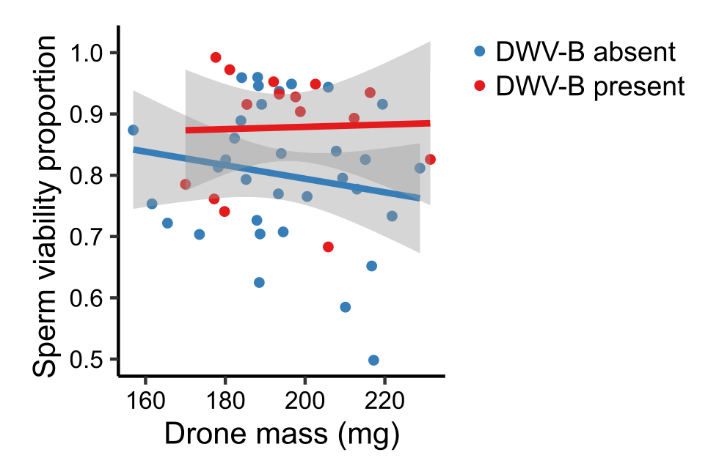

**Supplemental figure 1. Relationship between DWV-B presence and sperm viability.** Among untreated semen samples (time = 0), there was no significant relationship between drone mass and sperm viability (F = 0.65, df = 1, p = 0.43), but presence of underlying DWV-B infections was associated with higher sperm viability (F = 5.97, df = 1, p = 0.019) (linear model of arcsine square root transformed sperm viability against drone mass (continuous), and DWV-B (categorical, levels: presence, absence)).
